# Emergence of a novel mobile RND-type efflux pump gene cluster, *tmexC3D2-toprJ1b*, in *Pseudomonas* species

**DOI:** 10.1101/2021.07.26.453812

**Authors:** Shotaro Maehana, Ryotaro Eda, Nagi Niida, Aki Hirabayashi, Kouji Sakai, Takashi Furukawa, Kazunari Sei, Hidero Kitasato, Masato Suzuki

## Abstract

Tigecycline exhibits promising activity against multidrug-resistant Gram-negative bacteria (MDR-GNB), whose infections are difficult to treat with antimicrobials. However, mobile tigecycline resistance genes, such as *tmexCD-toprJ*, have emerged in Enterobacterales isolated in China, Vietnam, and possibly other countries in the world. In this study, we investigated tigecycline-nonsusceptible GNB in Japan. Eight tigecycline- and carbapenem-nonsusceptible isolates of *Pseudomonas alcaligenes* were obtained from sewage water from a medical institution in Japan in 2020. Whole genome analysis of all *P. alcaligenes* isolates was performed using short-read sequencing, and the isolate KAM426 was further analyzed using long-read sequencing. For important antimicrobial resistance genes, analysis of surrounding structures and comparison with similar sequences in the public genome database were performed. We identified a novel hybrid type of *tmexCD-toprJ* gene cluster, *tmexC3D2-toprJ1b* consisting of *tmexC3*, *tmexC2*, and *toprJ1b*, in phylogenetically clonal isolates of *P. alcaligenes*. The complete genome sequence of KAM426 revealed that this isolate co-harbors *tmexC3D2-toprJ1b* and two copies of the carbapenemase gene *bla*_IMP-1_ on the chromosome. *tmexC3D2-toprJ1b* in KAM426 was flanked by the IS*5*/IS*1182* family transposase gene, suggesting that the gene cluster was acquired by horizontal gene transfer (HGT). *tmexC3D2-toprJ1b* seemed to have spread to other *Pseudomonas* species such as *Pseudomonas aeruginosa* via HGT mediated by mobile gene elements such as a plasmid. This study identified *tmexCD-toprJ*-like tigecycline resistance genes in Japan for the first time and suggests that diverse *tmexCD-toprJ*-like gene clusters, including *tmexC3D2-toprJ1b*, have spread among MDR-GNB worldwide. Further epidemiological genomic studies in clinical and environmental settings are needed.

## Introduction

Tigecycline is considered a last-resort antimicrobial against infections caused by MDR Gram-negative bacteria. Recently, mobile tigecycline resistance genes, *tet*(X3), *tet*(X4), and other variants, *tet*(X5) to *tet*(X15), encoding flavin-dependent monooxygenases that catalyze tigecycline degradation have emerged in Enterobacterales and *Acinetobacter* species in China and other countries.^1–4^ Furthermore, mobile tigecycline resistance gene clusters, *tmexCD1-toprJ1*, *tmexCD2-toprJ2*, and *tmexCD3-toprJ1b* (also designated as *tmexCD3-toprJ3*), encoding the resistance–nodulation–cell division (RND) efflux pumps that excrete multiple antimicrobials, including tetracyclines such as tigecycline, cephalosporins, fluoroquinolones, and aminoglycosides, have emerged predominantly in Enterobacterales in China and Vietnam.^5–11^ Mobile *tmexCD-toprJ* genes are estimated to originate from chromosomal *mexCD-oprJ* genes in *Pseudomonas* species.^5^ Interestingly, however, *tmexCD1-toprJ1* has also been shown to spread to *Pseudomonas putida* by a megaplasmid.^7^

Here, we report *Pseudomonas alcaligenes* isolates harboring a novel variant of *tmexCD-toprJ* along with two copies of a metallo-β-lactamase (MBL) gene, *bla*_IMP-1_. *P. alcaligenes* is a Gram-negative aerobic rod belonging to the bacterial family Pseudomonadaceae, of which members are common inhabitants of soil and water and are rare opportunistic human pathogens.^12^ *P. alcaligenes* has also been suggested to be a causative agent of secondary bacterial infection during COVID-19 pneumonia.^13^ However, little is known about the clinical importance of *P. alcaligenes*, mainly because of the difficulties in identifying and distinguishing this bacterium from closely related *Pseudomonas* species such as *P. aeruginosa*, *P. mendocina,* and *P. pseudoalcaligenes*, in clinical settings.

## Results and discussion

Eight cephalosporin-resistant isolates of *P. alcaligenes* were obtained from medical wastewater in Japan in 2020. Whole-genome sequence analysis of *P. alcaligenes* isolates by Illumina HiSeq X and the core genome phylogeny based on their draft genome sequences showed that these isolates were phylogenetically very similar (Fig. S1). Moreover, all *P. alcaligenes* isolates harbored the same set of antimicrobial resistance (AMR) genes, including *tmexCD-toprJ*-like genes, *bla*_IMP-1_ (MBL gene conferring carbapenem resistance), *aac(6’)-Ib-cr* (aminoglycoside resistance gene), *fosE* (fosfomycin resistance gene), *qacG2* (multidrug resistance gene), and *sul1* (sulfonamide resistance gene), suggesting that these isolates were clonally disseminated (Fig. S1).

One of the *P. alcaligenes* isolates, KAM426, was further sequenced using ONT MinION, and hybrid sequence analysis using both Illumina and ONT reads resulted in the circular complete chromosome sequence (4.68 Mb, accession no. <u>AP024354</u>). Average nucleotide identity (ANI) analysis revealed that KAM426 is 96.5% identical to *P. alcaligenes* strain NCTC 10367^T^ (type strain, accession no. <u>UGUP00000000</u>), and the isolate harbored *tmexCD-toprJ*-like genes along with two copies of *bla*_IMP-1_ on its chromosome (Fig. 1). Also, *P. alcaligenes* KAM426 harbored a set of T4SS- and T6SS-associated genes, respectively, that would be potential virulence factors (Fig. 1). AST showed that *P. alcaligenes* KAM426 was nonsusceptible to tigecycline and broad-spectrum β-lactams, including carbapenems. According to the broth dilution method, the minimal inhibitory concentration (MIC) of tigecycline against KAM426 was 2 mg/L and was decreased to 1 mg/L in the presence of the efflux pump inhibitor NMP. According to the Etest, the MICs of imipenem and meropenem against KAM426 were 8 and >32 mg/L, respectively, and these were decreased to <1 and 0.19 mg/L, respectively, in the presence of the MBL inhibitor EDTA. These results suggested that *tmexCD-toprJ*-like genes and *bla*_IMP-1_ would be responsible for tigecycline and carbapenem resistance in this isolate. In *P. aeruginosa*, chromosomal efflux pump genes such as *mexCD-oprJ* (putative ancestor genes of *tmexCD-toprJ*) are known to contribute to tigecycline resistance.^5^ Though *P. alcaligenes* KAM426 did not harbor *mexCD-oprJ*-like genes on the chromosome, other chromosomal efflux pump genes could contribute tigecycline resistance in addition to *tmexCD-toprJ*-like genes.

**Fig. 1.**
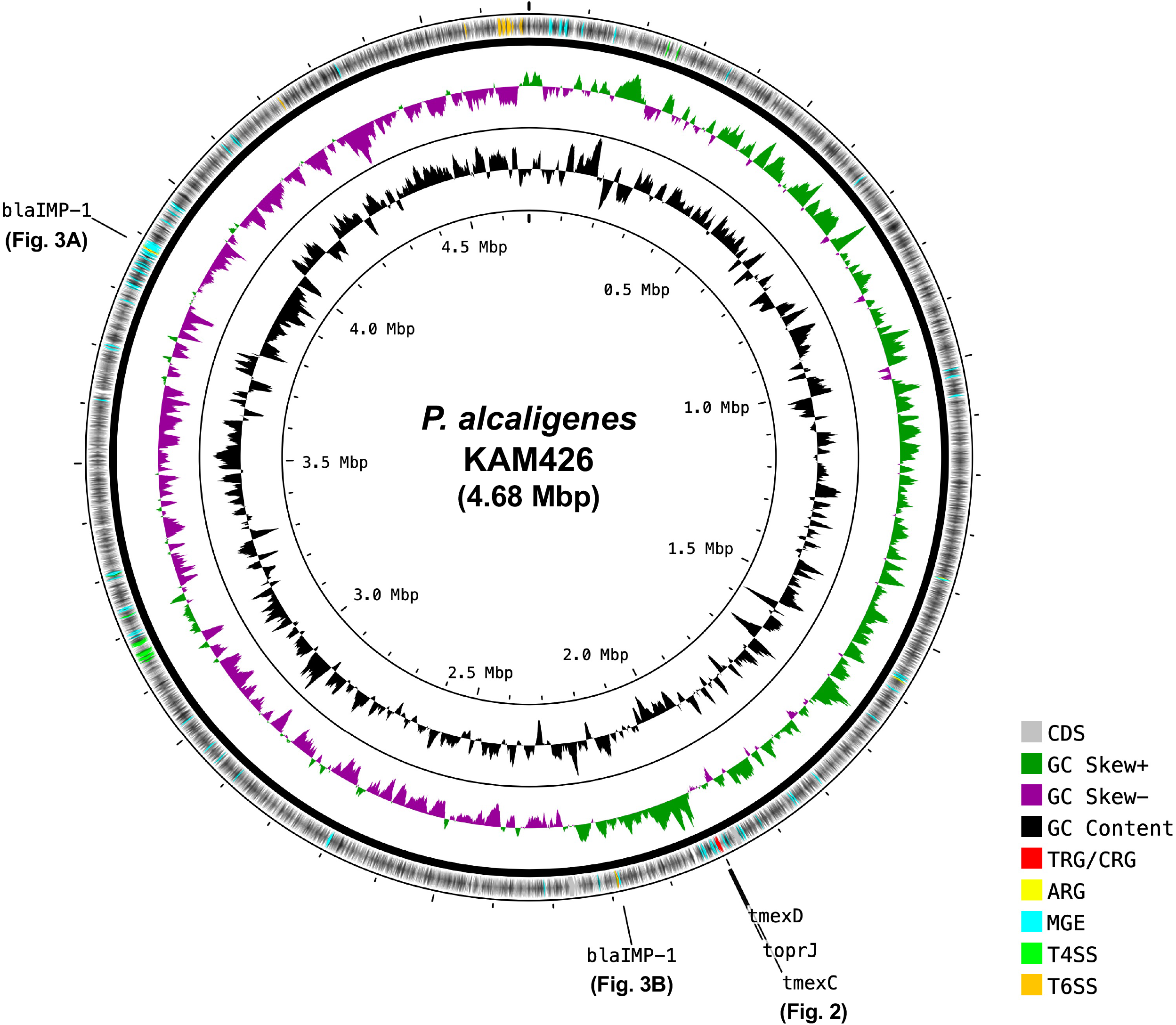
Circular representation of the chromosome of *Pseudomonas alcaligenes* KAM426 (accession no. <u>AP024354</u>) harboring *tmexC3D2-toprJ1b*, along with two copies of *bla*_IMP-1_ (shown in Figs. 2 and 3), isolated in Japan in 2020. Gray, green, purple, black, red, yellow, cyan, light green, and orange indicate coding sequences (CDS), GC skew+, GC skew-, GC content, tigecycline or carbapenem resistance genes (TRG/CRG), other AMR genes (ARG), mobile gene elements (MGE), type IV secretion system (T4SS)-associated genes, and type VI secretion system (T6SS)-associated genes, respectively.

The coding sequence of the *tmexCD-toprJ*-like gene cluster in *P. alcaligenes* KAM426 (1,999,524 to 2,005,275 nt in accession no. <u>AP024354</u>) was highly identical to that of *tmexCD1-toprJ1* in *Klebsiella pneumoniae* strain AH58I [96.8% (5,569/5,753 nt), 70,998 to 76,749 nt in accession no. <u>MK347425</u>] isolated from livestock in China in 2017,^5^ that of *tmexCD2-toprJ2* in *Raoultella ornithinolytica* strain NC189 [98.4% (5,660/5,752 nt), 182,964 to 188,715 nt in accession no. <u>MN175502</u>] isolated from a human in China in 2018,^8^ and that of *tmexCD3-toprJ1b* in *Proteus terrae* subsp. cibarius strain SDQ8C180-2T [98.1% (5,644/5,753 nt), 3,321,781 to 3,327,532 nt in accession no. <u>CP073356</u>] isolated from a chicken in China in 2018,^11^ respectively. The identities of the *tmexC*-like gene in KAM426 (KAM426_19240) compared with *tmexC1*, *tmexC2*, and *tmexC3* were 94.0% (1,094/1,164 nt), 94.6% (1,101/1,164 nt), and 98.5% (1,147/1,164 nt, the gene product was 98.4% identical to TMexC3 with six amino acid substitutions), respectively. For the *tmexD*-like gene in KAM426 (KAM426_19250), these identities were 96.5% (3,025/3,136 nt), 99.1% (3,108/3,135 nt, the gene product was perfect match to TMexD2), and 97.1% (3,046/3,136 nt), respectively. For the *toprJ*-like gene in KAM426 (KAM426_19260), the identities were 99.9% (1,433/1,434 nt), 99.9% (1,417/1,419 nt), and 100% (1,434/1,434 nt, the gene product was perfect match to TOprJ1b), respectively. Thus, we designated *tmexCD-toprJ*-like genes in KAM426 as *tmexC3D2-toprJ1b*.

The *tnfxB*-like gene, which has been suggested to be involved in the expression of the *tmexCD-toprJ*-like gene,^5^ was found upstream *tmexC3D2-toprJ1b* in *P. alcaligenes* KAM426 (KAM426_19230 in accession no. <u>AP024354</u>). The identities of the *tnfxB*-like gene in KAM426 compared with *tnfxB1*, *tnfxB2*, and *tnfxB3* (accession nos. <u>MK347425</u>, <u>CP054471</u>, and <u>CP073356</u>) were 97.7% (545/558 nt), 97.8% (540/552 nt), and 99.8% (557/558 nt, the gene product was 99.5% identical to TNfxB3 with one amino acid substitution), respectively. Thus, we designated the *tnfxB*-like gene in KAM426 as *tnfxB3*. *tnfxB3-tmexC3D2-toprJ1b* was flanked by the IS*5*/IS*1182* family transposase gene (Fig. 2A upper). Furthermore, the genomic region containing *tnfxB3*-*tmexC3D2-toprJ1b* in KAM426 was surrounded by many putative mobile gene elements (MGEs), and this genomic region was not present in *P. alcaligenes* strain NEB 585 (accession no. <u>CP014784</u>) (Fig. 2A). NEB 585 was isolated from a water environment in the United States in 1989 and is the only other *P. alcaligenes* strain for which the complete chromosome sequence has been reported^14^ other than KAM426. The genomic recombination regions in KAM426 and NEB 585 encoded a set of common genes (KAM426_19430 to KAM426_19470 in accession no. <u>AP024354</u>, and A0T30_13575 to A0T30_13555 in accession no. <u>CP014784</u>) (Fig. 2A), although their functions are unknown. The results suggest that KAM426 acquired *tmexCD-toprJ*-like genes, which confer resistance to multiple antimicrobials including tigecycline, via horizontal gene transfer (HGT) mediated by MGEs.

**Fig. 2.**
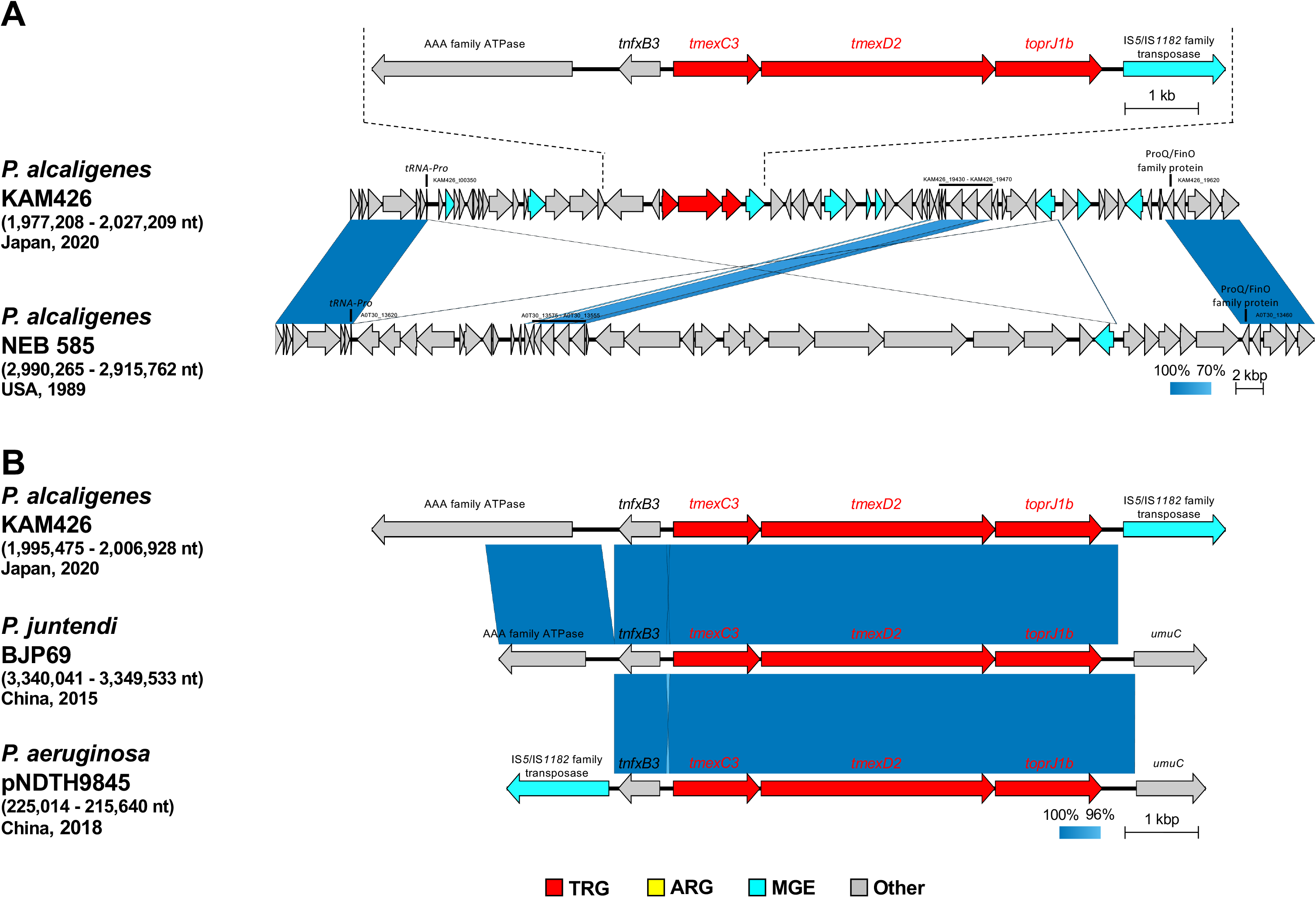
The *tmexC3D2-toprJ1b* gene cluster in *Pseudomonas alcaligenes* KAM426. (A) Genetic context of the *tmexC3D2-toprJ1b* gene cluster in *P. alcaligenes* KAM426 and its surrounding genomic region (the region between 1,977,208 and 2,027,209 nt in accession no. <u>AP024354</u>), and structural comparison with the corresponding genomic region in *P. alcaligenes* NEB 585 (the region between 2,990,265 and 2,915,762 nt in accession no. <u>CP014784</u>). (B) Structural comparison of the *tmexC3D2-toprJ1b* gene cluster in *P. alcaligenes* KAM426 (the region between 1,995,475 and 2,006,928 nt in accession no. <u>AP024354</u>) with that in the chromosome of *P. juntendi* BJP69 (the region between 3,340,041 and 3,349,533 nt in accession no. <u>CP041933</u>) and in plasmid pNDTH9845 of *P. aeruginosa* NDTH9845 (the region between 225,014 and 215,640 nt in accession no. <u>CP073081</u>). The strain names of *Pseudomonas* species, along with the country and year in which bacteria were isolated, are shown. *tmexC3D2-toprJ1b* genes (TRG), other AMR genes (ARG), mobile gene elements (MGE), and other genes (Other) are highlighted in red, yellow, light blue, and gray, respectively. Sequence identity is shown as a color scale with the indicated percentages.

BLASTn analysis using megablast revealed that two *Pseudomonas* species strains in the NCBI database of Nucleotide collection (nr/nt) have the exact same sequence containing the *tmexC3D2-toprJ1b* gene cluster along with *nfxB*. A *bla*_DIM-2_-harboring *Pseudomonas* species strain, BJP69, isolated from a human in China in 2015^15^ carries *tmexC3D2-toprJ1b* on its chromosome (accession no. <u>CP041933</u>) and a *bla*_KPC-2_-harboring *P. aeruginosa* strain, NDTH9845, isolated from a human in China in 2018 carries *tmexC3D2-toprJ1b* on its IncP-2 megaplasmid, pNDTH9845 (accession no. <u>CP073081</u>) (Fig. 2B). ANI analysis confirmed that BJP69 is 98.0% identical to *Pseudomonas juntendi* strain BML3^T^ (type strain, accession no. <u>BLJG01000000</u>) and that NDTH9845 is 99.2% identical to *P. aeruginosa* DSM 50071^T^ (type strain, accession no. <u>FUXR01000000</u>). *tmexC3D2-toprJ1b* was determined to be flanked by the IS*5*/IS*1182* family transposase gene in *P. aeruginosa* pNDTH9845, whereas no MGE was found upstream or downstream of *tmexC3D2-toprJ1b* in *P. juntendi* BJP69 (Fig. 2B). Together with *P. alcaligenes* KAM426 in this study, these three *Pseudomonas* species strains harbor acquired carbapenemase genes, in addition to *tmexC3D2-toprJ1b*, showing that they have accumulated clinically relevant AMR genes. Of note, *P. juntendi* BJP69 carried one more copy of *tmexCD-toprJ*-like genes, *tmexC2D2-toprJ2*, on its IncP-2 megaplasmid, pBJP69-DIM (accession no. <u>MN208064</u>).

The integron-integrase IntI1 catalyzes site-specific recombination between the *attI1* and *attC* sites.^16^ The class 1 integron gene cassette consisting of *intI1* with the *attI1* site, *qacE∆1* (disrupted form of *qacE*), and *sul1* in *P. alcaligenes* KAM426 (accession no. <u>AP024354</u>) contain several AMR genes, including *fosE*, two copies of *aac(6’)-Ib-cr*, *bla*_IMP-1_, and *qacG2* with their *attC* sites (Fig. 3A). The *bla*_IMP-1_-containing integron gene cassette in KAM426 was found to be surrounded by many putative transposase genes, and this MGE-containing genomic region was not present in *P. alcaligenes* NEB 585 (accession no. <u>CP014784</u>) (Fig. 3A). The genomic region around ATP-dependent helicase genes (KAM426_37950 and KAM426_36180 in accession no. <u>AP024354</u>, and A0T30_05350 in accession no. <u>CP014784</u>) could be a hot spot for HGT (Fig. 3A), but there have been no reports to suggest this possibility to date.

**Fig. 3.**
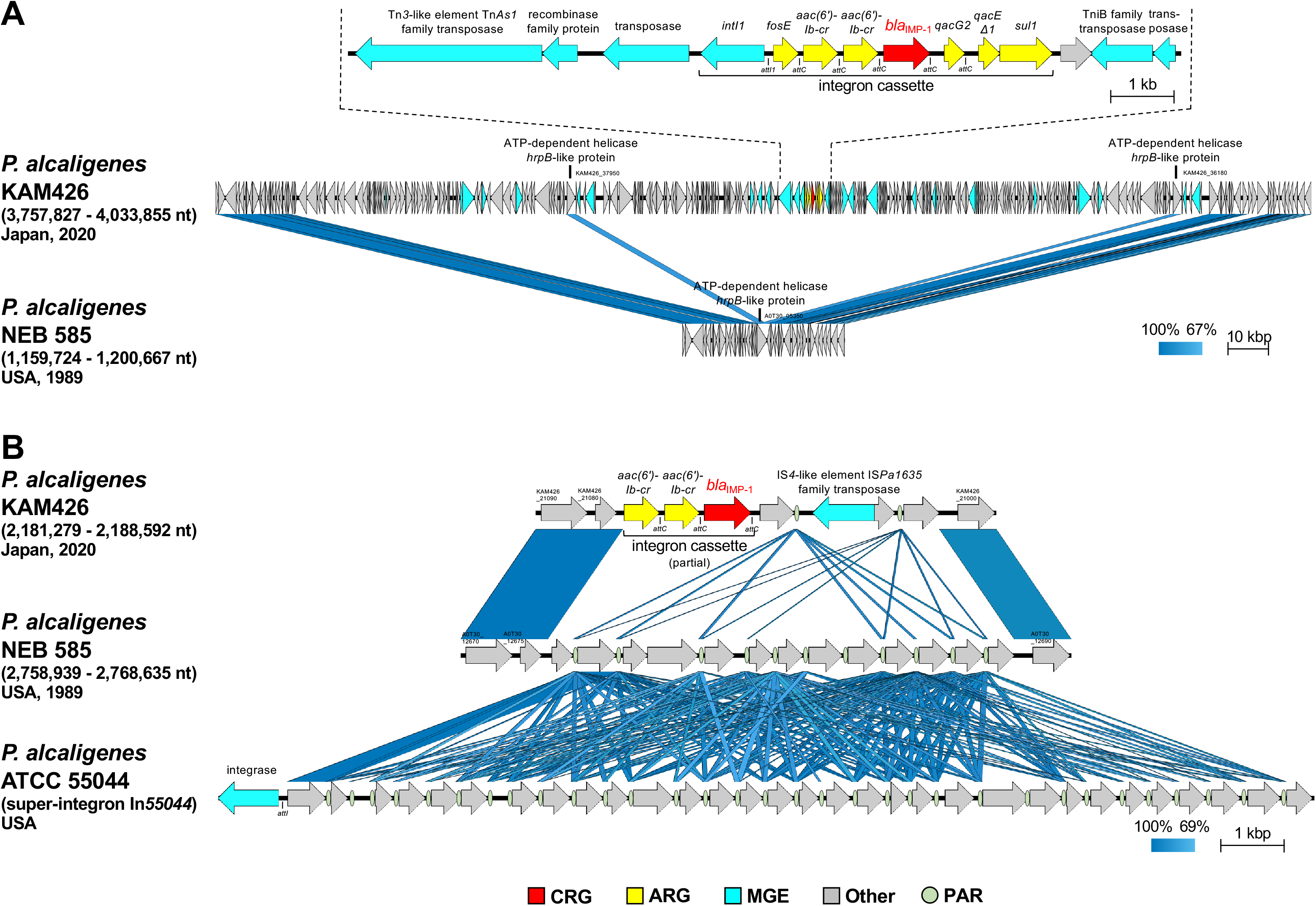
Two copies of *bla*_IMP-1_ genes in *Pseudomonas alcaligenes* KAM426. (A) Genetic context of the class I integron gene cassette containing *bla*_IMP-1_ in *P. alcaligenes* KAM426 and its surrounding genomic region (the region between 3,757,827 and 4,033,855 nt in accession no. <u>AP024354</u>), and structural comparison with the corresponding genomic region in *P. alcaligenes* NEB 585 (the region between 1,159,724 and 1,200,667 nt in accession no. <u>CP014784</u>). (B) Genetic context of the partial integron gene cassette containing the other *bla*_IMP-1_ gene in *P. alcaligenes* KAM426 and its surrounding genomic region (the region between 2,181,279 and 2,188,592 nt in accession no. <u>AP024354</u>), and structural comparison with the corresponding genomic regions in *P. alcaligenes* NEB 585 (the region between 2,758,939 and 2,768,635 nt in accession no. <u>CP014784</u>) and in *P. alcaligenes* ATCC 55044 (super-integron In*55044* in accession no. <u>AY038186</u>). The strain names of *P. alcaligenes*, along with the country and year in which bacteria were isolated, are shown. *bla*_IMP-1_ genes (CRG), other AMR genes (ARG), mobile gene elements (MGE), other genes (Others), and *P. alcaligenes* repetitive DNA (PAR) are highlighted in red, yellow, light blue, gray, and khaki green, respectively. Sequence identity is shown as a color scale with the indicated percentages.

The other copy of *bla*_IMP-1_ was contained within a partial structure of the integron gene cassette consisting of two copies of *aac(6’)-Ib-cr* followed by *bla*_IMP-1,_ as described previously herein (Fig. 3A), in a different location in the chromosome of *P. alcaligenes* KAM426 (Fig. 3B). Interestingly, a comparison between the *bla*_IMP-1_-containing genomic region of *P. alcaligenes* KAM426 and the corresponding genomic region in *P. alcaligenes* NEB 585 revealed the presence of multiple copies of short repeated sequences (approximately 80 bp), which were identical to PARs (*<u>P</u>seudomonas <u>a</u>lcaligenes* <u>r</u>epetitive <u>D</u>NAs) within the super-integron In*55044* in *P. alcaligenes* strain ATCC 55044 (accession no. <u>AY038186</u>)^17^ (Fig. 3B).

The super-integron, which was first identified in *Vibrio cholerae*, is distinguished from conventional integrons in several respects, such as size and the nature of the genes contained within cassettes and contributes to the acquisition of AMR genes.^18–20^ The PARs were reported as recombination sites for the integrase gene (*intI*_Pac_ in accession no. <u>AY038186</u>) in the super-integron in *P. alcaligenes*.^17^ The PARs in *P. alcaligenes* KAM426 and *P. alcaligenes* NEB 585 contained conserved sequences of inverted repeats (1L, 2L, 2R, and 1R) with a PAR signature and variable regions between inverted repeats 2L and 2R, as shown in *P. alcaligenes* ATCC 55044^17^ (Fig. S2). Although KAM426 and NEB 585 lacked the integrase gene flanking the PAR and most of the contained genes had no known function, these plastic genomic regions in both strains retained their evolutionary histories of gene acquisitions mediated by the super-integron. There was no PAR around *bla*_IMP-1_ in KAM426, suggesting that the super-integron is not directly involved in the acquisition of *bla*_IMP-1_, and this genomic region is likely one of the hot spots for HGT. KAM426 is thought to have incorporated *bla*_IMP-1_ into the class 1 integron gene cassette first (Fig. 3A) and then incorporated the partial structure containing *bla*_IMP-1_ into the other genomic region flanked by the super-integron (Fig. 3B), leading to a high level of resistance by increasing the copy number of AMR genes.

Our study provides a glimpse into environmental bacteria that have been rapidly and silently becoming resistant to clinically relevant antimicrobials, including tigecycline and carbapenem, and highlights the importance of AMR monitoring using wastewater to detect a future clinical crisis before it happens. Furthermore, *P. alcaligenes* would be considered an important environmental reservoir that supplies AMR genes to other related *Pseudomonas* species that are more virulent and likely to cause nosocomial infections, such as *P. aeruginosa*.

## Materials and methods

### Bacterial isolation and antimicrobial susceptibility testing

Eight cephalosporin-resistant isolates of *P. alcaligenes* (KAM426, KAM428, KAM429, KAM430, KAM432, KAM434, KAM435, and KAM436) were obtained from sewage water from a medical institution in Japan in February, 2020. Environmental water samples were collected and cultured using DHL (Deoxycholate Hydrogen sulfide Lactose) agar containing 2 mg/L of ceftriaxone. Bacterial species identification was performed using MALDI Biotyper (Bruker). antimicrobial susceptibility testing (AST) using *Escherichia coli* ATCC 25922 as quality control was performed according to the broth dilution method based on the CLSI 2020 guidelines or according to the Etest (bioMérieux) based on the manufacturer instructions. For tigecycline, AST was additionally performed in the presence or absence of 75 mg/L of the efflux pump inhibitor 1-(1-naphthylmethyl)-piperazine (NMP) as used in the previous study.^5^

### Whole-genome sequencing and subsequent bioinformatics analysis

Whole-genome sequencing of all eight cephalosporin-resistant isolates of *P. alcaligenes* was performed using HiSeq X (Illumina), and the isolate KAM426 was further sequenced using MinION [Oxford Nanopore Technologies (ONT)] with the R9.4.1 flow cell. The library for Illumina sequencing (paired-end, insert size of 500-900 bp) was prepared using Nextera XT DNA Library Prep Kit and the library for ONT sequencing was prepared using Rapid Barcoding Kit (SQK-RBK004). Illumina reads were assembled de novo using Shovill v1.1.0 (https://github.com/tseemann/shovill) with default parameters, resulting in the draft genome sequences. For KAM426, ONT reads were basecalled using Guppy v4.2.2 with the high-accuracy mode, and then both Illumina and ONT reads were assembled de novo using Unicycler v0.4.8.0 (https://github.com/rrwick/Unicycler) with default parameters, resulting in the complete circular chromosome sequence (accession no.: <u>AP024354</u>).

Coding sequence (CDS) annotation and average nucleotide identity (ANI) analysis were performed using the DFAST server (https://dfast.nig.ac.jp). Antimicrobial resistance (AMR) genes were detected using ResFinder v4.1 (http://www.genomicepidemiology.org) with default parameters using the customized AMR gene database including all known *tmexCD-toprJ* genes. Type IV secretion system (T4SS)- and type VI secretion system (T6SS)-associated genes were detected using TXSScan v1.0.5 (https://research.pasteur.fr/en/tool/txsscan-models-and-profiles-for-protein-secretion-systems/). Circular genomic sequence was visualized using the CGView server (http://cgview.ca). Linear comparison of sequence alignment was performed using BLAST and visualized using Easyfig v.2.2.2 (http://mjsull.github.io/Easyfig/).

### Nucleotide Sequences

The complete genome sequence of *P. alcaligenes* KAM426 has been deposited at GenBank/EMBL/DDBJ under the accession number <u>AP024354</u>. Draft genome sequences of *P. alcaligenes* KAM428, KAM429, KAM430, KAM432, KAM434, KAM435, and KAM436 have been deposited at GenBank/EMBL/DDBJ under the accession numbers <u>BPMN00000000</u>, <u>BPMO00000000</u>, <u>BPMP00000000</u>, <u>BPMQ00000000</u>, <u>BPMR00000000</u>, <u>BPMS00000000</u>, and <u>BPMT00000000</u>, respectively.

## Acknowledgments

This work was supported by grants (JP21fk0108093, JP21fk0108139, JP21fk0108133, JP21wm0325003, JP21wm0325022, JP21wm0225004, JP21wm0225008, and JP21gm1610003 to M. Suzuki) from the Japan Agency for Medical Research and Development (AMED), and grants (20K07509 to M. Suzuki; 21K15440 to A. Hirabayashi) from the Ministry of Education, Culture, Sports, Science and Technology (MEXT), Japan.

---

Fig. S1. Core genome phylogeny constructed by Roary v3.13.0 (https://github.com/sanger-pathogens/Roary) with minimum percentage identity for BLASTp=70% and RAxML v8.2.4 (https://github.com/stamatak/standard-RAxML), with 1,000 bootstraps using ceftriaxone-resistant *Pseudomonas alcaligenes* isolates in this study and reference strains of *P. alcaligenes* and *P. aeruginosa* (NCTC 10367^T^ and NEB 585 for *P. alcaligenes*, and PAO1 for *P. aeruginosa*). *P. aeruginosa* was used as the outgroup. Bar lengths represent the number of substitutions per site in the core genome. Detected AMR genes, genome assembly status (complete or draft genome sequence, contig numbers if draft), sizes, and accession numbers are shown.

Fig. S2. Alignment of *Pseudomonas alcaligenes* repetitive DNAs (PARs) in *P. alcaligenes* strains. One PAR in *P. alcaligenes* ATCC 55044 (super-integron In*55044* in accession no. <u>AY038186</u>), two PARs in *P. alcaligenes* KAM426 (accession no. <u>AP024354</u>), and 11 PARs in *P. alcaligenes* NEB 585 (accession no. <u>CP014784</u>) are shown. The multiple alignment comparison was performed and visualized using MAFFT v7 (https://mafft.cbrc.jp/alignment/software/). The PAR signature sequence and variable region are shown. Open boxes and arrows represent consensus sequences of inverted repeats (1L, 2L, 2R, and 1R), as described previously.^17^

