## Supplemental Figs. 1 and 2 for "Emergence of a novel mobile RND-type efflux pump gene cluster, *tmexC3D2-toprJ1b*, in *Pseudomonas* species"

### Fig. S1

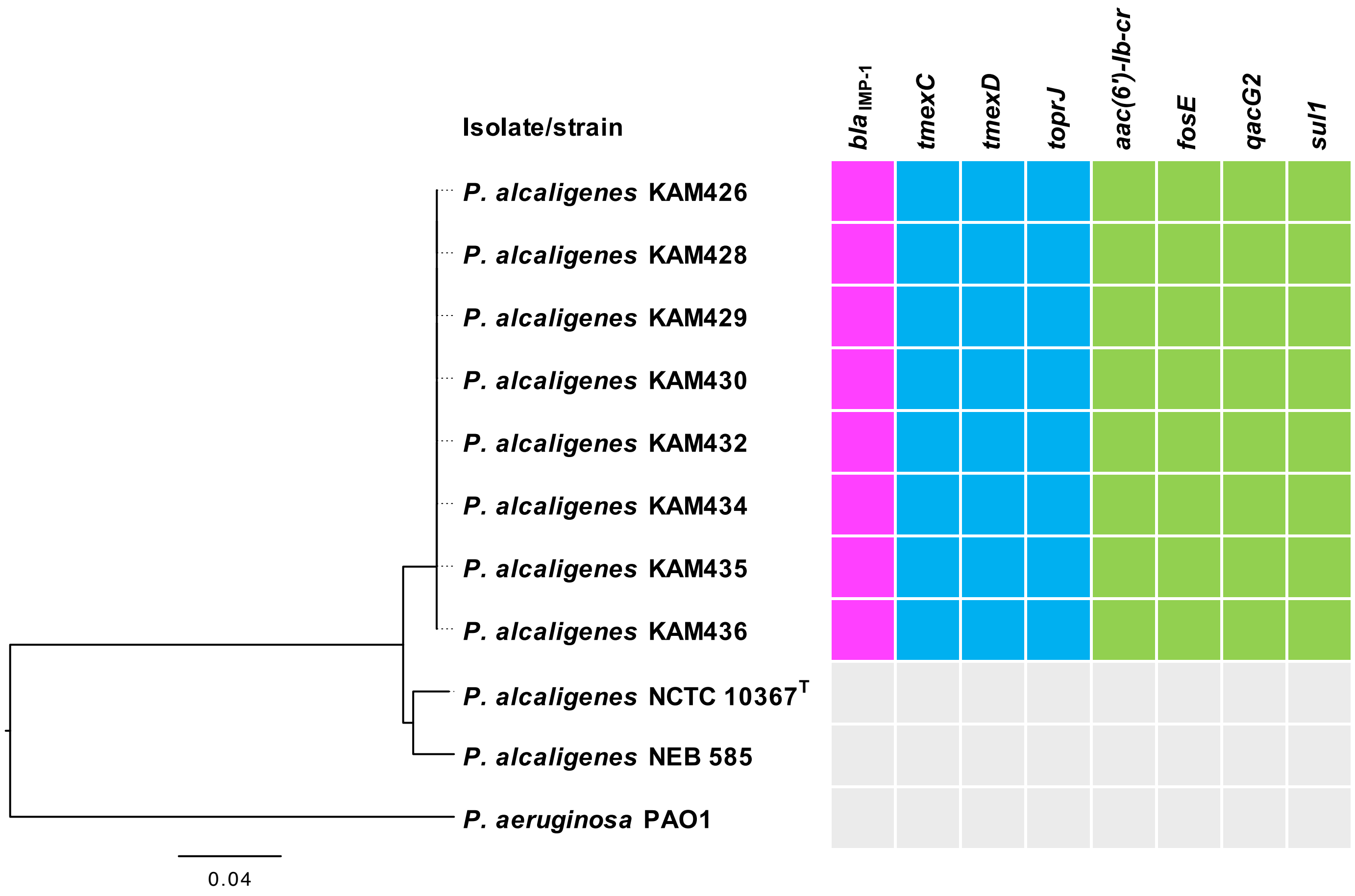


| Isolate/strain | Genome | Size (bp) | Accession. no. |
|---|---|---|---|
| *P. alcaligenes* KAM426 | Complete | 4,680,436 | AP024354 |
| *P. alcaligenes* KAM428 | Draft (104 contigs) | 4,511,893 | BPMN00000000 |
| *P. alcaligenes* KAM429 | Draft (105 contigs) | 4,543,302 | BPMO00000000 |
| *P. alcaligenes* KAM430 | Draft (93 contigs) | 4,537,340 | BPMP00000000 |
| *P. alcaligenes* KAM432 | Draft (99 contigs) | 4,539,395 | BPMQ00000000 |
| *P. alcaligenes* KAM434 | Draft (93 contigs) | 4,541,115 | BPMR00000000 |
| *P. alcaligenes* KAM435 | Draft (97 contigs) | 4,540,209 | BPMS00000000 |
| *P. alcaligenes* KAM436 | Draft (98 contigs) | 4,538,303 | BPMT00000000 |
| *P. alcaligenes* NCTC 10367$^T$ | Draft (2 contigs) | 5,016,483 | UGUP00000000 |
| *P. alcaligenes* NEB 585 | Complete | 4,406,305 | CP014784 |
| *P. aeruginosa* PAO1 | Complete | 6,217,899 | AE004091 |

### Fig. S2

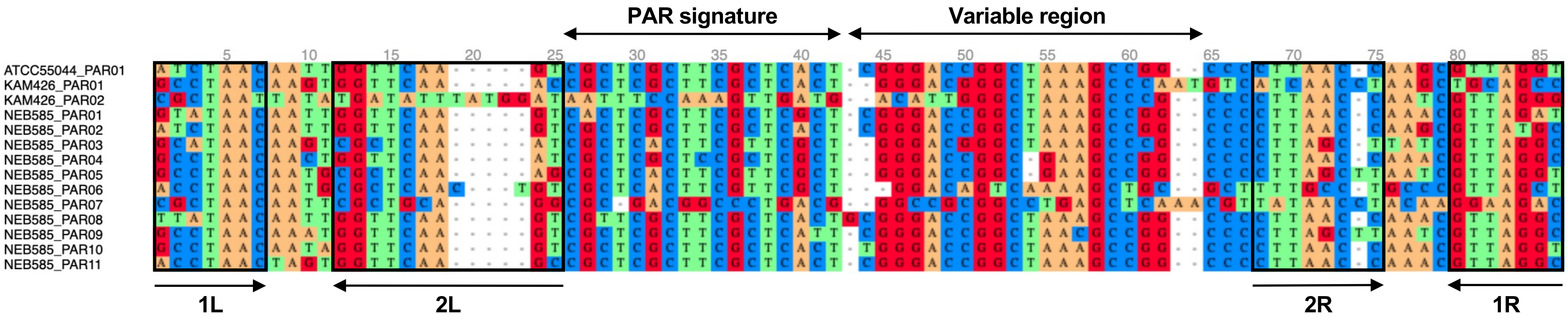
PAR signature
Variable region
ATCC55044_PAR01
KAM426_PAR01
KAM426_PAR02
NEB585_PAR01
NEB585_PAR02
NEB585_PAR03
NEB585_PAR04
NEB585_PAR05
NEB585_PAR06
NEB585_PAR07
NEB585_PAR08
NEB585_PAR09
NEB585_PAR10
NEB585_PAR11
1L
2L
2R
1R
